## Supplemental Material for "An isoform-specific function of Cdc42 in regulating mammalian Exo70 during axon formation"

### Figure S1

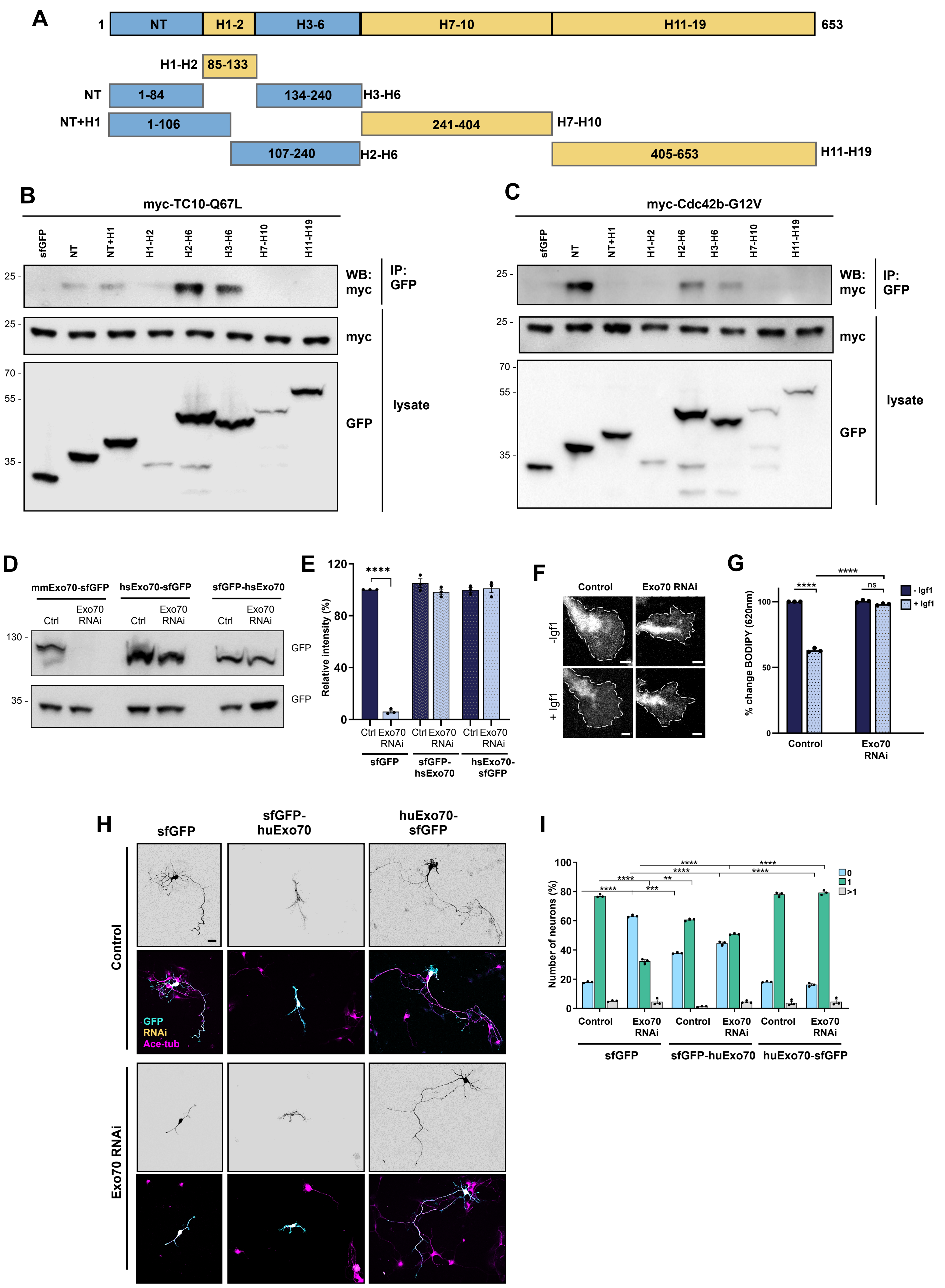

A
1
NT
H1-2
H3-6
H7-10
H11-19
653
H1-H2
85-133
NT
1-84
134-240
H3-H6
NT+H1
1-106
241-404
H7-H10
107-240
H2-H6
405-653
H11-H19
B
myc-TC10-Q67L
sfGFP
NT
NT+H1
H1-H2
H2-H6
H3-H6
H7-H10
H11-H19
WB: myc
IP: GFP
myc
lysate
GFP
C
myc-Cdc42b-G12V
D
mmExo70-sfGFP
hsExo70-sfGFP
sfGFP-hsExo70
Ctrl
Exo70 RNAi
GFP
E
Relative intensity (%)
****
sfGFP
sfGFP-hsExo70
hsExo70-sfGFP
F
Control
Exo70 RNAi
-Igf1
+ Igf1
G
% change BODIPY (620nm)
- Igf1
+ Igf1
ns
H
sfGFP
sfGFP-huExo70
huExo70-sfGFP
Control
Exo70 RNAi
GFP
RNAi
Ace-tub
I
Number of neurons (%)
0
1
>1
***
**

**Figure S1. Interaction of TC10 and Cdc42palm with Exo70**

(A) Schematic representation of the Exo70 fragments that were used for interaction studies. The numbers indicate the position of amino acids in the coding sequence.

(B and C) HEK 293T cells were co-transfected with vectors for sfGFP-Exo70-NT (aa 1-84), -NT+H1 (aa 1-106), -H1-H2 (aa 85-106), -H2-H6 (aa 107-240), -H3-H6 (aa 133-240), -H7-H10 (aa 241-404), or -H11-H19 (aa 405-653) and myc-TC10-Q67L (B) or myc-Cdc42b-G12V (C), immunoprecipitated with an anti-GFP nanobody (IP) and bound proteins analyzed by Western blot (WB). The molecular weight is indicated in kDa.

(D) Validation of the Exo70 knockdown vector. HEK 293T cells were co-transfected with vectors for mouse Exo70-sfGFP (mmExo70-sfGFP) or human Exo70 tagged at the N-terminus (sfGFP-hsExo70) or C-terminus (hsExo70-sfGFP) and control or knockdown vectors for Exo70 expressing EmGFP. The efficiency of the Exo70 knockdown was analyzed by Western blot using an anti-GFP antibody. Human Exo70 is resistant to the knockdown vector due to mismatches in the nucleotide sequence.

(E) Quantification of the knockdown efficiency from (D).

(F) The knockdown of Exo70 blocks Igf1-induced exocytosis. Hippocampal neurons from E18 rat embryos were transfected with control vector or knockdown vectors for Exo70 expressing H2B-mRFP. Neurons were incubated with BODIPY FL-C5 ceramide at 1 d.i.v. to label post-Golgi vesicles and cultured in medium without (-Igf1) or with 20 nM Igf1 (+Igf1) for 10 minutes. Igf1-stimulated exocytosis results in a decrease in the fluorescence signal for BODIPY-labeled vesicles in growth cones at 620 nm. Scale bars: 2 µm.

(G) Igf1-stimulated exocytosis was quantified as the decrease in the fluorescence intensity for BODIPY-labeled vesicles in growth cones at 620 nm normalized to the signal at 515 nm. The relative change in fluorescence intensities for BODIPY-labeled vesicles is shown for (F). The value for control neurons (-Igf1) was set to 100% (one-way ANOVA followed by Tukey's test, N=27 neurons from each condition from n=3 independent experiments).

(H) Expression of hsExo70-sfGFP but not sfGFP-hsExo70 rescues the knockdown of Exo70 in neurons. Hippocampal neurons from E18 rat embryos were co-transfected with control or knockdown vectors for Exo70 expressing H2B-RFP (yellow) and vectors for sfGFP-hsExo70 or hsExo70-sfGFP (cyan). Neurons were analyzed at 3 d.i.v. with an anti-acetylated tubulin antibody (magenta). Scale bar: 20 µm.

(I) The percentage of neurons without an axon (0), with a single axon (1) and with multiple axons (>1) was quantified from (H) (unpaired Student's t-test, n=3 independent experiments, N= 250 neurons). Interestingly, a fusion of sfGFP to the N-terminus of Exo70 (sfGFP-Exo70), which is similar to N-terminally tagged constructs that were used in some previous studies (Lira et al., 2018; Liu et al., 2007; Zuo et al., 2006) did not rescue the knockdown.

Values are means ± SEM.

### Figure S2

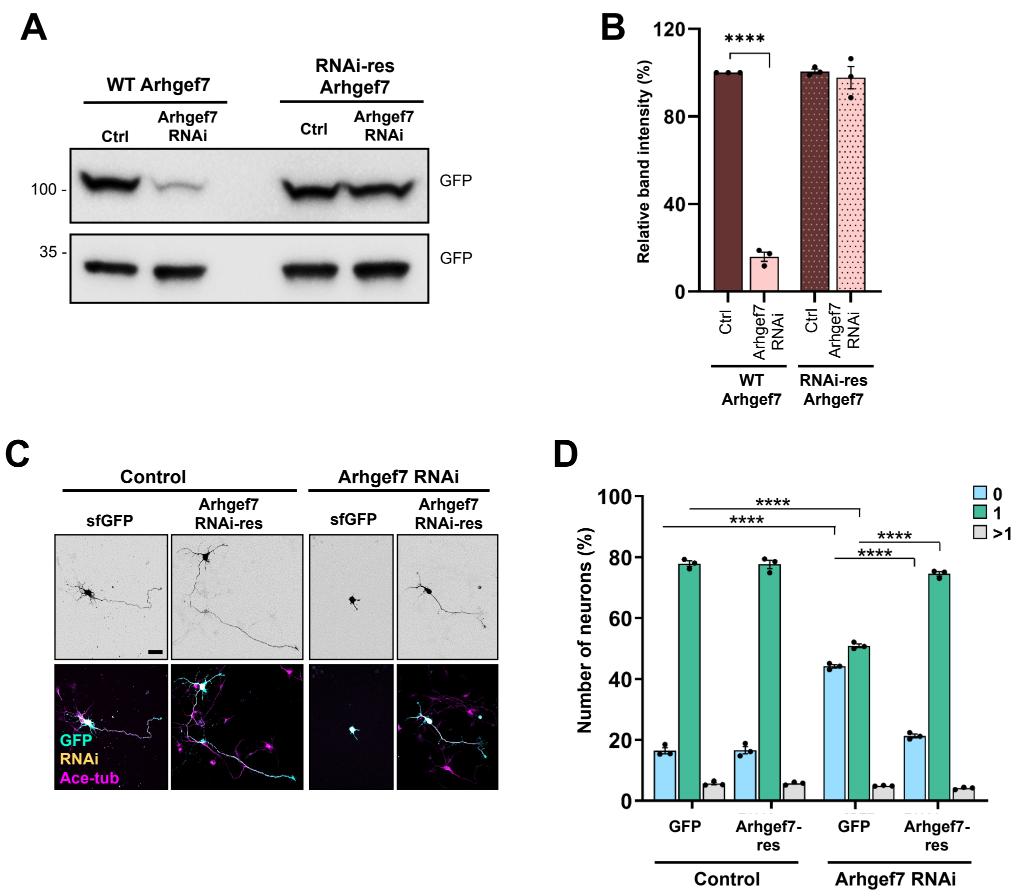
A
WT Arhgef7
RNAi-res Arhgef7
Ctrl
Arhgef7 RNAi
Ctrl
Arhgef7 RNAi
100 -
35 -
GFP
GFP
B
Relative band intensity (%)
120
80
40
0
****
Ctrl
Arhgef7 RNAi
Ctrl
Arhgef7 RNAi
WT Arhgef7
RNAi-res Arhgef7
C
Control
Arhgef7 RNAi
sfGFP
Arhgef7 RNAi-res
sfGFP
Arhgef7 RNAi-res
GFP
RNAi
Ace-tub
D
Number of neurons (%)
100
80
60
40
20
0
****
****
****
****
0
1
>1
GFP
Arhgef7-res
GFP
Arhgef7-res
Control
Arhgef7 RNAi

**Figure S2. Validation of the knockdown vector for Arhgef7**

(A) HEK 293T cells were co-transfected with vectors for wild-type (Arhgef7) or RNAi-resistant sfGFP-Arhgef7 (RNAi-res Arhgef7) and control or knockdown vectors for Arhgef7 expressing EmGFP. The efficiency of the knockdown was analyzed by Western blot (WB) using an anti-GFP antibody.

(B) The efficiency of the Arhgef7 knockdown was quantified for (A) (unpaired Student's t-test, n=3 independent experiments).

(C) Hippocampal neurons from E18 rat embryos were co-transfected with control or knockdown vectors for Arhgef7 expressing H2B-RFP (yellow) and vectors expressing sfGFP or RNAi-resistant sfGFP-Arhgef7 (cyan). Neurons were analyzed at 3 d.i.v. with an anti-acetylated tubulin antibody (magenta). Scale bar: 20 µm.

(D) The percentage of neurons without an axon (0), with a single axon (1) and with multiple axons (>1) was quantified for (C) (unpaired Student's t-test, n=3 independent experiments, N= 301 neurons).

Values are means ± SEM.

### Figure S3

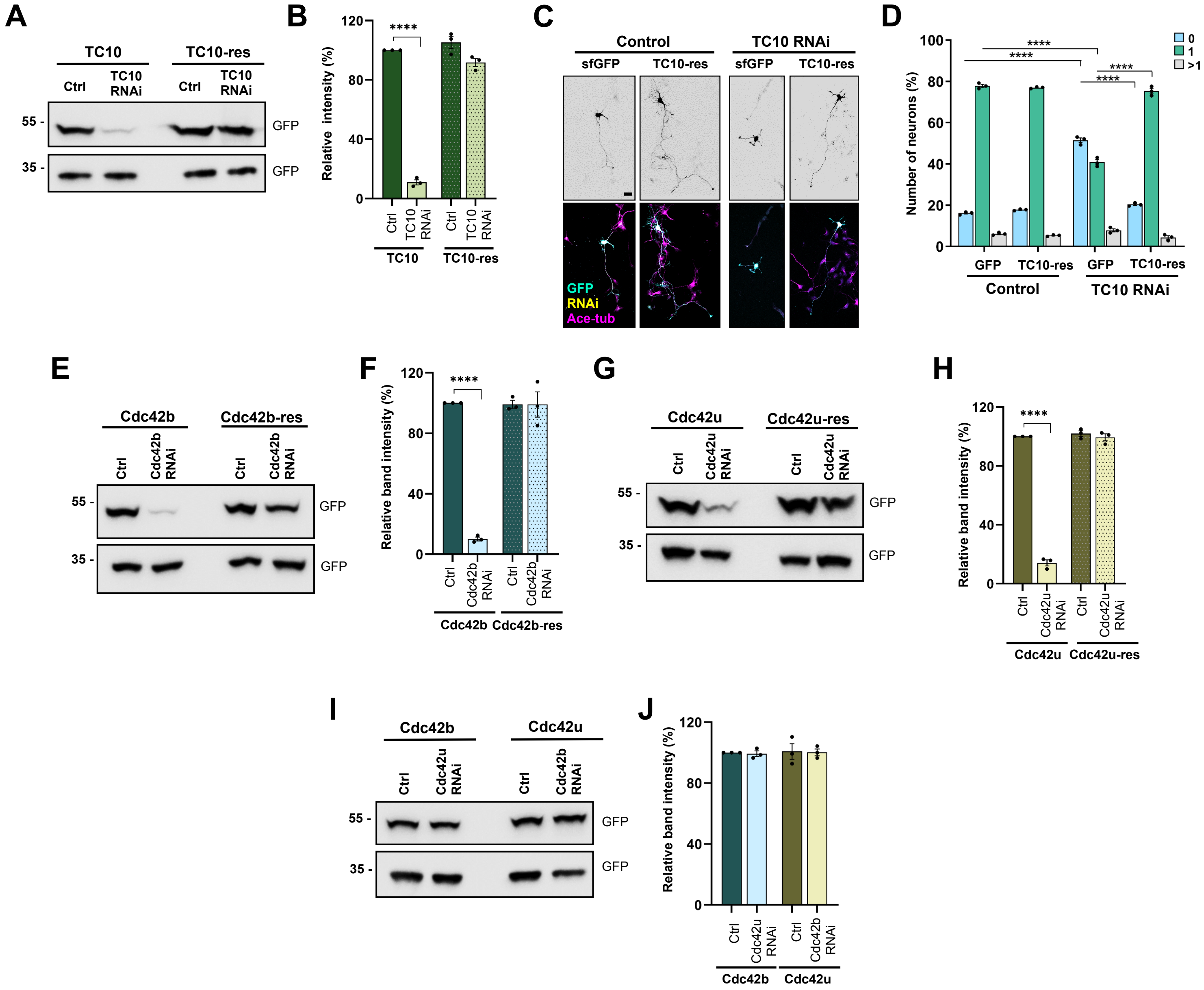
A
TC10
TC10-res
Ctrl
TC10 RNAi
55 -
35 -
GFP
B
Relative intensity (%)
****
TC10
TC10-res
C
Control
TC10 RNAi
sfGFP
TC10-res
GFP
RNAi
Ace-tub
D
Number of neurons (%)
0
1
>1
****
GFP
TC10-res
Control
TC10 RNAi
E
Cdc42b
Cdc42b-res
Ctrl
Cdc42b RNAi
F
Relative band intensity (%)
G
Cdc42u
Cdc42u-res
Cdc42u RNAi
H
I
J

**Figure S3. Validation of knockdown vectors for TC10, Cdc42b and Ccd42u**

(A) HEK 293T cells were co-transfected with vectors for wild-type (TC10) or RNAi-resistant sfGFP-TC10 (RNAi-res TC10) and control or knockdown vectors for TC10 expressing EmGFP. The efficiency of the knockdown was analyzed by Western blot (WB) using an anti-GFP antibody.

(B) The efficiency of the TC10 knockdown was quantified for (A) (n=3 independent experiments, unpaired Student's t-test).

(C) Hippocampal neurons from E18 rat embryos were co-transfected with control or knockdown vectors for TC10 expressing H2B-RFP (yellow) and vectors expressing sfGFP or RNAi-resistant sfGFP-TC10 (cyan). Neurons were analyzed at 3 d.i.v. with an anti-acetylated tubulin antibody (magenta). Scale bar: 20 μm.

(D) The percentage of neurons without an axon (0), with a single axon (1) and with multiple axons (>1) was quantified for (C) (unpaired Student's t-test, n=3 independent experiments, N=238 neurons).

(E) HEK 293T cells were co-transfected with vectors for wild-type (Cdc42b) or RNAi-resistant EGFP-Cdc42b (RNAi-res Cdc42b) and control or knockdown vectors for Cdc42b expressing EmGFP. The efficiency of the knockdown was analyzed by Western blot using an anti-GFP antibody.

(F) The efficiency of the Cdc42b knockdown was quantified for (E) (n=3 independent experiments, unpaired Student's t-test).

(G) HEK 293T cells were co-transfected with vectors for wild-type (Cdc42u) or RNAi-resistant EGFP-Cdc42u (RNAi-res Cdc42u) and control or knockdown vectors for Cdc42u expressing EmGFP. The efficiency of the knockdown was analyzed by Western blot using an anti-GFP antibody.

(H) The efficiency of the Cdc42u knockdown was quantified for (G) (n=3 independent experiments, unpaired Student's t-test).

(I) The isoform-specific knockdown vectors do not target the other Cdc42 isoform. HEK 293T cells were co-transfected with vectors for EGFP-Cdc42u or -Cdc42b and control or knockdown vectors expressing EmGFP as indicated. The efficiency of the knockdown was analyzed by Western blot using an anti-GFP antibody.

(J) The efficiency of the knockdown was quantified for (I) (n=3 independent experiments, unpaired Student's t-test).

Values are means ± SEM.

| Table S1 (Oligonuceotides), Related to STAR Methods - Plasmids | |
|---|---|
| **Primer name** | **Sequence 5’-3’** |
| sFGFP-C3 reverse | CAGATCTGAGTACTTGTACAGCTCGTCCATGC |
| mmExo70-sfGFP forward | CGGAATTCTGATGATTCCCCCGCAGGAGG |
| hsExo70-sfGFP forward | CGGAATTCTGATGATTCCCCCACAGGAGG |
| Exo70-sfGFP reverse | GCGTCGACACGGCAGAGGTGTCGAAAAGGCG |
| Exo70 NT reverse | GCGTCGACTTACAGGCAGGACAGCGTCTTCTC |
| Exo70 NT+H1 reverse | GCGTCGACTAATGTGGGGCCCTCTCTGATG |
| Exo70 H1-H2 forward | CGGAATTCTGATGGACCATGTCATCAGCTACTACC |
| Exo70 H1-H2 reverse | GCGTCGACTTAGTCTGGGCTGTTGTCCTGG |
| Exo70 H2-H6 forward | CGGAATTCTGATGGGTAGGCTGGAAGAGTACCTGG |
| Exo70 H3-H6 forward | GCGAATTCTGATGAGCCCGGAACTCAACAAAGTG |
| Exo70 H2-H6 reverse | GCGTCGACTAACTTATGGAAATGCTCCTTC |
| Exo70 H7-H10 forward | CGGAATTCTGATGAGCAGTTCTTCCTCTGGGG |
| Exo70 H7-H10 reverse | GCGTCGACTAAATTCTTGATGTTGTCTGC |
| Exo70 H11-H19 forward | CGGAATTCTGATGGACCCGGACAAGGAGTAC |
| Exo70 miRNA Top strand | TGCTGTCAAACAGCAGCTTCACTTTGGTTTTGGCCA CTGACTGACCAAAGTGACTGCTGTTTGA |
| Exo70 miRNA Bottom strand | CCTGTCAAACAGCAGTCACTTTGGTCAGTCAGTGGC CAAAACCAAAGTGAAGCTGCTGTTTGAC |
| Arhgef7 miRNA Top strand | TGCTGATAACCTTCAGGATCTGAGCGGTTTTGGCCA CTGACTGACCGCTCAGACTGAAGGTTAT |
| Arhgef7 miRNA Bottom strand | CCTGATAACCTTCAGTCTGAGCGGTCAGTCAGTGGC CAAAACCGCTCAGATCCTGAAGGTTATC |
| TC10 miRNA Top strand | TGCTGAACACAGTCTTCAAACCCTTCGTTTTGGCCAC TGACTGACGAAGGGTTAAGACTGTGTT |
| TC10 miRNA Bottom strand | CCTGAACACAGTCTTAACCCTTCGTCAGTCAGTGGC CAAAACGAAGGGTTTGAAGACTGTGTTC |
| Cdc42b miRNA Top strand | TGCTGTTTGGGTTGAGTTTCCGGAGGGTTTTGGCCA CTGACTGACCCTCCGGACTCAACCCAAA |
| Cdc42b miRNA Bottom strand | CCTGTTTGGGTTGAGTCCGGAGGGTCAGTCAGTGG CCAAAACCCTCCGGAAACTCAACCCAAAC |
| Cdc42u miRNA Top strand | TGCTGCACACCTGCGGCTCTTCTTCGGTTTTGGCCA CTGACTGACCGAAGAAGCCGCAGGTGTG |
| Cdc42u miRNA Bottom strand | CCTGCACACCTGCGGCTTCTTCGGTCAGTCAGTGGC CAAAACCGAAGAAGAGCCGCAGGTGTGC |
| Arhgef7 RNAi rescue forward | GCCCGTGTCACCCAACTCAGGGACATTGAAGAGCC CTC |
| Arhgef7 RNAi rescue reverse | GAGGGCTCTTCAATGTCCCTGAGTTGGGTGACACGG GC |
| TC10 RNAi rescue forward | GCTTTAACCCAGAAGGGCTTGAACACTGTGTTTGAT GAGG |
| TC10 RNAi rescue reverse | CCTCATCAAACACAGTGTTCAAGCCCTTCTGGGTTAA AGC |
| Cdc42b/u forward | CGGGATCCATGCAGACAATTAAGTGTG |
| Cdc4b reverse | TTGCGGCCGCTTAGAATATACTGCTCTTCCTTTTGGG TTGAGTTTCCGGAGGCTCGAGGGC |
| Cdc42b RNAi rescue reverse | CCGCGGCCGCTTAGAATATACAGCACTTCCTTTTGG GCTGGGTCTCCGGAGGCTCGAGGGC |
| Cdc42u RNAi rescue reverse | TTGCGGCCGCTCATAGCAGCACACACCTGCGACTGT TGTTCGGTTCTGG |
